# Spatiotemporal changes in biodiversity by ecosystem engineers: how beavers structure the richness of large mammals

**DOI:** 10.1101/2020.12.02.406785

**Authors:** Lindsay Y. Gauvin, Daniel Gallant, Eric Tremblay, Dominique Berteaux, Nicolas Lecomte

**Author notes:** Senior author (Nicolas Lecomte). Current addresses: Kouchibouguac National Park of Canada, Kouchibouguac, New Brunswick, E4X 2P1, Canada (Daniel Gallant).

## Abstract

High levels of biodiversity may be needed to maintain ecosystem functioning. By creating niches for other species, ecosystem engineers have the potential to promote biodiversity, but it is unclear how this translates across spatiotemporal scales. We evaluated the long-term impact of ecosystem engineering by beavers (*Castor canadensis*) on the diversity of mobile species. We tested the hypothesis that the spatial distribution of engineered habitats in different states resulting from ecosystem engineering by beavers increases the biodiversity of large mammals across spatial scales. We tested a second hypothesis that engineered habitat in different states resulting from ecosystem engineering by beavers drive the diversity of large mammals. We compared the richness and composition of boreal mammals using camera traps between habitats with and without history of occurrence by beavers within a protected area, where trapping, hunting, and forest exploitation are prohibited. We found that unique species were mostly found in specific engineered habitats, with ponds and wet meadows showing more species than dry meadows. In addition to the increased diversity of dispersal-limited species, our results show that beavers promote the diversity of mobile species at both local and landscape scales, signaling the importance of niche creation in structuring animal communities across scales.

## Introduction

Current changes in global biodiversity^1^ have sparked interest in various aspects of its relationship with ecosystem functioning, one of them being the link between biodiversity and the stable supply of ecosystem goods and services. Recent studies suggest that high levels of biodiversity may be needed to maintain ecosystem functioning^2,3,4,5^. Much of the relationship between biodiversity and ecosystem functioning may be determined by the diversity of species’ traits^6^. The ability of some species to create niches for other species (e.g. ecosystem engineering)^7^ seems to be a trait that has a disproportionate weight on the functioning of many ecosystems, by building or modifying habitat structure^8^. Although studies demonstrated that ecosystem engineers can promote higher species richness^9,10,11,12^, few of them have quantified this potential on various spatiotemporal scales^13^. Our understanding of the impacts of ecosystem engineers on food webs is therefore still at an early stage^14,15,16,17^.

Beavers (*Castor canadensis*) are textbook examples of ecosystem engineers because their stream impoundment activities and foraging habits strongly modify their surrounding environment, which then influences plant and animal communities^18^. Dam building by beavers produces wetlands, which when abandoned by beavers, transition into meadows that can persist for more than 50 years^19,20^. Overall, this beaver-modified habitat process leads to a mosaic of heterogeneous habitats in different states across the landscape^21^. A recent meta-analysis focused on published studies measuring and illustrating the impact of beavers on invertebrates, plants, amphibians, reptiles, birds and mammals^22^. Overall, such meta-analysis signalled how much beavers, by creating habitats, have a highly positive influence on biodiversity. Yet it is not clear whether ecosystem engineering by beavers influences species richness at local scales. For instance, in several studies, beaver wetlands increased species richness for reptiles^23,24^, while positive effects^25,26,27^, negative effects^24^ or no effect^23^ on species diversity and abundance were reported for amphibians. However, one study demonstrated that study scale was critical when assessing the impact of beavers on plant communities^13^. At the local scale, the effect of engineered patches on herbaceous plant species richness was measured as neutral. However, there were high vs. low levels of species similarity and richness among non-engineered vs. engineered patches, respectively^13^. Thus, species richness was higher when combining non-engineered and engineered patches, showing a positive effect of engineered patches on species richness at the landscape scale^13^. Nevertheless, how such biodiversity shifts occur over the longterm is still poorly understood. In addition, past studies focused on dispersal-limited species, as well as those with small home ranges^25,26,27^. Recent studies show that habitats created by beavers had a positive influence on local species richness of several types of mobile species; however, these species were also highly associated to wetlands or to open habitats in general (e.g. ducks^28,29^ and bats^30,31^). Mobile species are characterized by individuals with regular and predictable movements (migrants) to erratic resource-driven movements (nomads). All large mammals in our study area are considered nomad species with movements ranging from less than a kilometer (e.g. Raccoon) to about ten kilometers (e.g. Moose) within their respective home ranges. While early reports demonstrated that some large mammals benefit from beaver ponds and meadows^32^, we know little about the scale at which these mobile species with larger home ranges may respond to habitat patches created by beavers (but see^33^).

We studied the long-term impact of ecosystem engineering by beavers on the diversity of large mammals at both local and landscape scales. Here diversity is a generic term and to measure it we quantified species richness and composition. In addition, local scale is defined as the local habitat patch (engineered or non-engineered). The effect at the landscape scale was tested by comparing the combined biodiversity of engineered and non-engineered patches to strictly engineered patches and to strictly non-engineered patches. We tested the hypothesis (Hyp. 1) that the spatial distribution of engineered habitats in different states resulting from foraging and dam building by beavers increases the diversity of large mammals across spatial scales. We predicted that engineered patches host a higher frequency of occurrence for species found in both patch types than non-engineered patches (Prediction 1.1), along with higher species richness at the local and landscape scales (Prediction 1.2). Here occurrence refers to an event (i.e. detection of an animal) and frequency of occurrence is defined as the number of times the event occurred during the length of our study. We tested a second hypothesis (Hyp. 2), that engineered habitat in different states resulting from ecosystem engineering by beavers drive the diversity of large mammals. We predicted differences in the frequency of species occurrence between ponds and meadows (Prediction 2). To test these predictions, we used automatic cameras to compare the richness and composition of boreal mammals between habitats with and without a history of beaver occurrence within a protected area, where trapping, hunting, and forest exploitation are prohibited.

## Results

A total of 16 979 images triggered by movement were visually examined by one person to reduce observer bias. Images were visually examined three times and number of detections did not vary among examinations. There were 125 single detections within a 24-hour period with identifiable animals comprising 9 mammalian species and 8 detections with unidentifiable animals captured by camera traps from June 6^th^ to August 26^th^ in 2014 and May 27^th^ to August 18^th^ in 2015. This amounts to a total of 1,132 camera days at 16 sites belonging to multiple watersheds of Kouchibouguac National Park of Canada (see Supplementary Fig. S1 online). We tested the effect of year on the frequency of species occurrence and species richness and it was not retained in our models. The 9 mammalian species were moose (*Alces alces*; 33 detections), white-tailed deer (*Odocoileus virginianus*; 29 detections), black bear (*Ursus americanus*; 43 detections), raccoon (*Procyon lotor*; 3 detections), coyote (*Canis latrans*; 4 detections), red fox (*Vulpes vulpes*; 1 detection), bobcat (*Felis rufus*; 1 detection), snowshoe hare (*Lepus americanus*; 10 detections) and muskrat (*Ondatra zibethicus*; 1 detection) (see Supplementary Table S1). We did not consider beavers in our analyses although they were only detected by cameras at beaver sites.

### Ecosystem engineering and frequency of species occurrence

The frequency of species occurrence between engineered and non-engineered patches and the generalized linear model demonstrated that ecosystem engineering by beavers has an effect on the frequency of mammalian species occurrence (Prediction 1.1, Fig.1; see Supplementary Table S2). Engineered and non-engineered patches shared four species (black bear, white-tailed deer, moose, and raccoon). However, the frequency of occurrence of individual species demonstrated that moose were three times as frequent at engineered patches compared to non-engineered patches (0.044 ± 0.012, CI: 0.021–0.067 vs. 0.014 ± 0.006, CI: 0.003–0.026, Fig. 1). Black bear was, on average, twice as frequent at engineered patches compared to non-engineered patches (0.052 ± 0.011, CI: 0.031–0.074 vs. 0.025 ± 0.012, CI: 0.002–0.049), Fig. 1). However, the difference was not statistically significant because the confidence interval overlap was more than 25% due to high variance within patch types. Also, four species were found exclusively at engineered patches (bobcat, coyote, red fox, and muskrat), whereas snowshoe hare was found exclusively at non-engineered patches.

**Figure 1:**
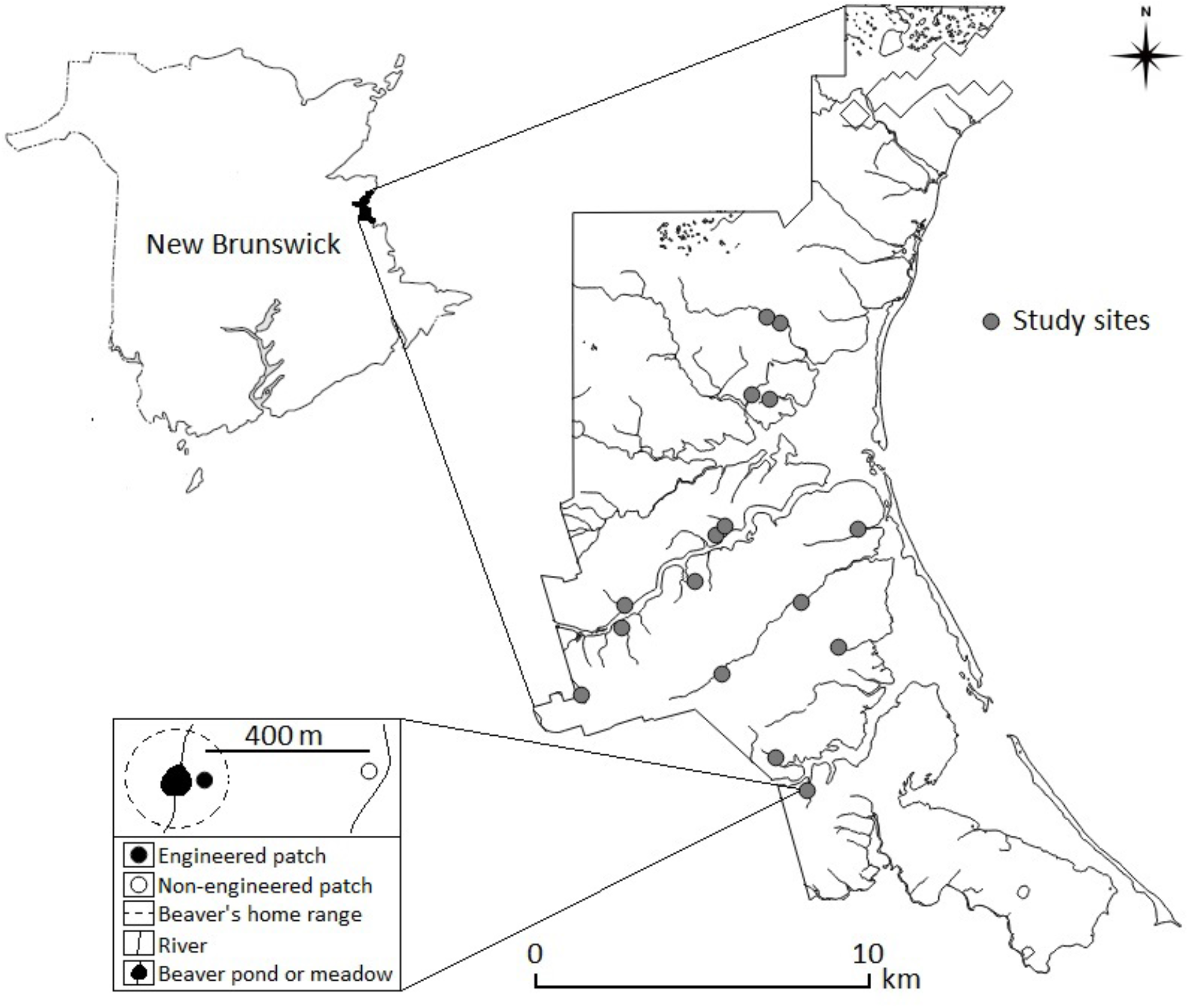
Frequency of species occurrence showing the effect of ecosystem engineering by beavers on the frequency of occurrence of individual species at **(a)** non-engineered and **(b)** engineered patches (both n= 16) in Kouchibouguac National Park of Canada, 2014–2015. We divided the total number of detections of each species by the number of camera trap days at each patch and calculated the mean and the standard error for both patch types. The detection period for each patch was over a 6-month period (i.e. 3 months per year). Error bars represent + 1 SE. Animal silhouettes were acquired from an open source (http://openclipart.org/).

### Ecosystem engineering and species composition

The ordination analysis demonstrated that ecosystem engineering by beavers has an effect on the species composition of mobile mammalian species (Prediction 1.2); species composition is similar among engineered patches (i.e., patches located close to each other in the ordination), and dissimilar among non-engineered patches (i.e., patches dispersed in the ordination) (Fig. 2A). In addition, moose, coyote, and black bear were close together, hence showing they were often documented together at individual patches. The same situation occurred for snowshoe hare and raccoon, as well as for raccoon and bobcats, while muskrats, red foxes, and white-tailed deer were not associated with any other species. Also, Fig. 2A shows that most engineered and non-engineered patches within each study site were separated (e.g. compare the location of E1 to that of U1), indicating that our data does not suffer from double or multiple detections of the same individuals within study sites. Study sites with the same location (3 out of 16) were a result of low species frequencies (i.e. the same one or two species). Patches without any detection were excluded from the ordination because they had no ordination score (U8 and U9).

**Figure 2:**
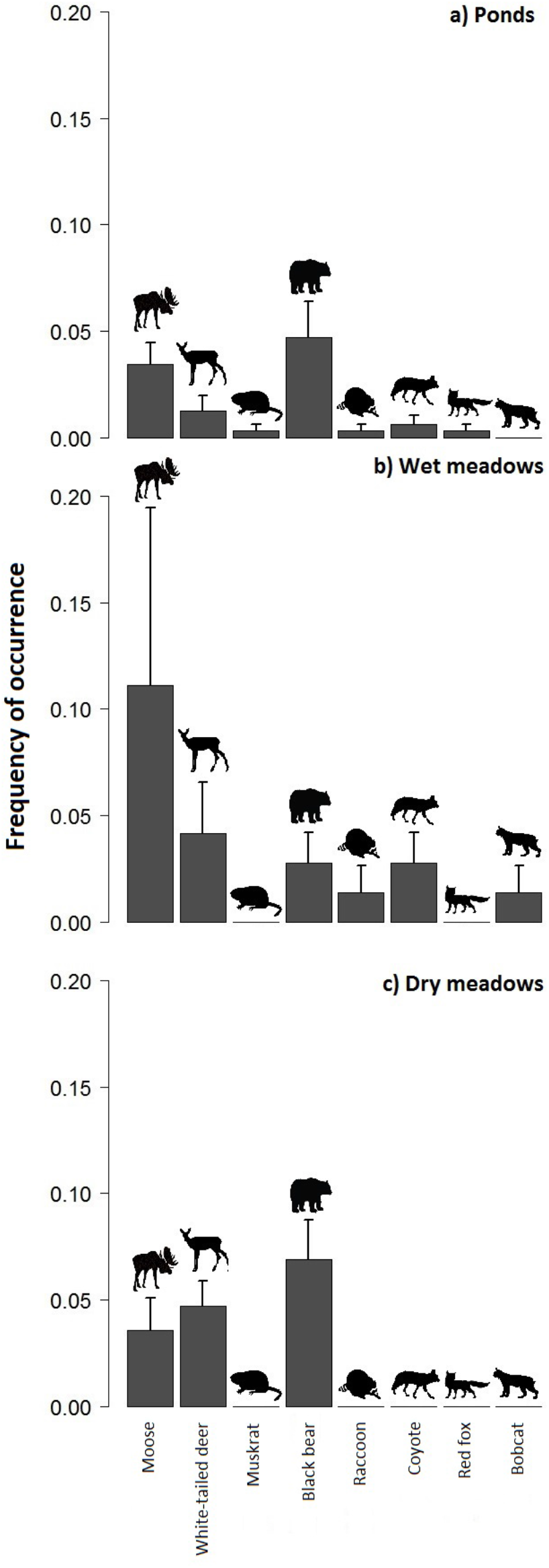
Effect of ecosystem engineering by beavers on species composition: **(a)** ordination of animal community composition between engineered and non-engineered patches (both n=16), Kouchibouguac National Park of Canada, 2014–2015. Ordination of patch based on presence of species using non-metric multidimensional scaling. Species and patches are situated according to their similarity to other species or patches; the closer they are to each other in the ordination the more similar they are. Numbers represent patch ID, (E) engineered patch, (U) non-engineered patch, and black hollow circles represent the exact location of the patch ID linked by a black line. Patches without any detection were excluded from the ordination as they can’t be given a score (i.e. U8 and U9). Silhouettes represent (from top to bottom): snowshoe hare, raccoon, bobcat, white-tailed deer, moose, coyote, black bear, red fox, and muskrat (silhouette centers represent exact locations). Animal silhouettes were acquired from an open source (http://openclipart.org/). **(b)** Similarity in community composition within and between non-engineered and engineered patches, and **(c)** within and between ponds, wet meadows, and dry meadows. Mean Morisita-Horn similarity index for pairwise comparisons of patches of the same and different type based on the frequency of occurrence of species. Error bars represent + 1 SE. Bars with distinct letters are significantly different.

The mean Morisita-Horn similarity indices demonstrated that ecosystem engineering by beavers has an effect on the species composition of mobile mammalian species (Prediction 1.2). Non-engineered patches were more variable in composition than engineered patches (0.32 ± 0.04, CI: 0.25–0.39 vs. 0.55 ± 0.03, CI: 0.50–0.60, Fig. 2B). Comparisons of patches from engineered and non-engineered habitat indicate an intermediate level of species overlap between the two habitat types (0.41 ± 0.02, CI: 0.37–0.45). In total, 44% of the 9 mammalian species recorded in the survey were found in both engineered and non-engineered patches.

### Ecosystem engineering and species richness

Species richness of engineered and non-engineered patches and the generalized linear mixed model demonstrated that ecosystem engineering by beavers has an effect on the richness of mobile mammalian species at a local scale (Prediction 1.2). Species richness was twice as high in individual engineered patches as compared to non-engineered patches (0.076 ± 0.009, CI: 0.058–0.094 vs. 0.038 ± 0.006, CI: 0.026–0.050; only a marginal CI overlap) (Fig. 3A, Table 1). The generalized linear mixed model demonstrated that patch type played a significant role on the observed species richness (p < 0.001). Estimated species richness and the linear model demonstrated that ecosystem engineering by beavers has an effect on the richness of mobile mammalian species at a landscape scale (Prediction 1.2). Species richness estimated for all combined engineered patches was marginally higher (11 ± 2.45, CI: 6–16) compared to all non-engineered patches (6 ± 1.41, CI: 3–9, Fig. 3B, Table 2) based on their CI. Compared to all engineered patches, species richness estimated for all study sites was, however, not higher (12 ± 2.45, CI: 7–17). The linear model demonstrated that patch type played a significant role in the observed estimated species richness (p < 0.001) while study sites did not (p = 0.939).

**Table 1:** Results of a generalized linear mixed model showing the effect of patch type (engineered vs. non-engineered) on total species richness at 16 study sites, Kouchibouguac National Park of Canada, 2014–2015.

| Coefficient | Coefficient estimate | Standard error | *Z* | *P* |
| --- | --- | --- | --- | --- |
| Species | 2.153 | 0.320 | 6.72 | <0.0001 |
| Patch type | -0.815 | 0.231 | -3.53 | <0.0001 |

Note: Variance and standard deviation for random factor patches within each study site was 4.39E-05±6.625E-03, respectively. Patch type is coded 1 (engineered) and 2 (non-engineered). Marginal $R^2 = 0.28$ .

**Table 2:** Results of a linear model showing the effect of patch type (engineered vs. non-engineered) on species richness estimated from jackknife estimators at 16 study sites, Kouchibouguac National Park of Canada, 2014–2015.

| Factors | Estimate | Standard error | *Z* | *P* |
| --- | --- | --- | --- | --- |
| Engineered patches | 11.616 | 0.768 | 15.129 | <0.0001 |
| Study sites | -0.075 | 0.978 | -0.076 | 0.939 |
| Non-engineered patches | -5.304 | 1.252 | -4.236 | <0.0001 |

Note: Adjusted $R^2 = 0.18$ .

**Figure 3:**
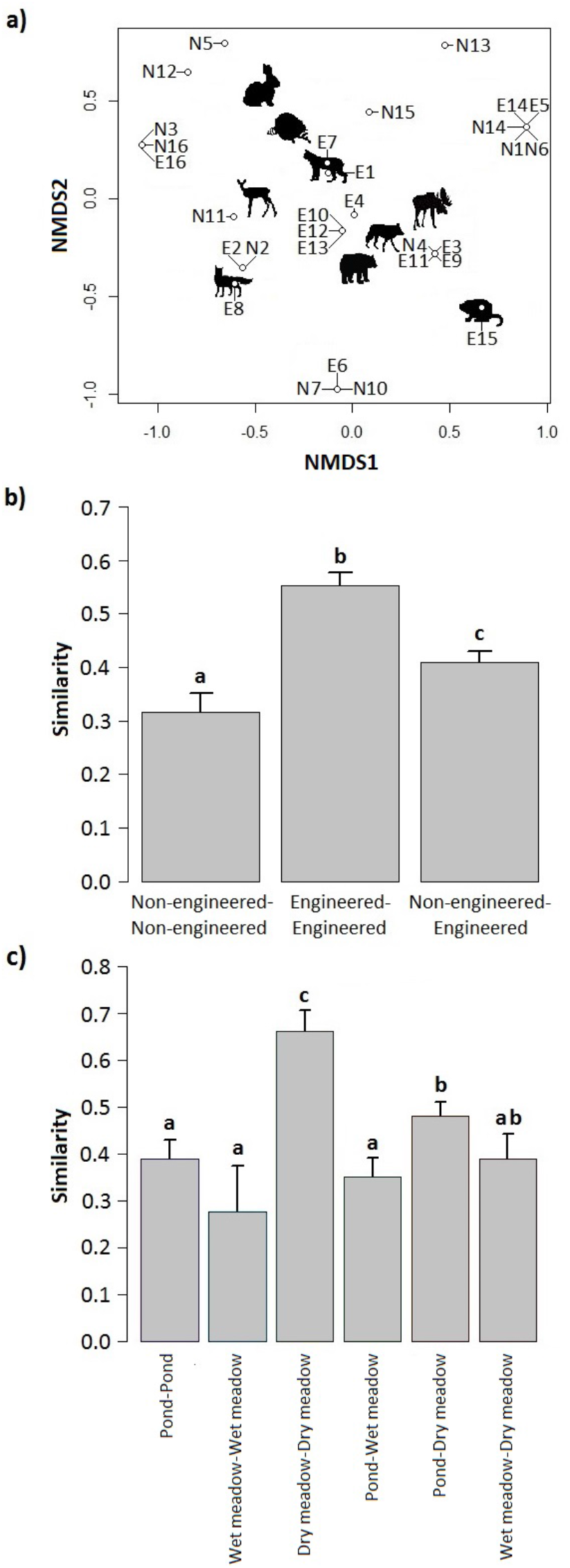
Effect of ecosystem engineering by beavers on species richness on multiple scales: **(a)** species richness of non-engineered patches and engineered patches, **(b)** species richness estimated applying the first order jackknife estimator from all non-engineered patches, all engineered patches, and all engineered and non-engineered patches, **(c)** species richness of ponds, wet meadows and dry meadows, and **(d)** species richness estimated applying the first order jackknife estimator from all ponds, all wet meadows, and all dry meadows, Kouchibouguac National Park of Canada, 2014–2015. We divided the number of species by the number of camera days at each patch and calculated the mean and the standard error for patch types. Error bars represent + 1 SE. Bars with distinct letters are significantly different.

### Habitat states and frequency of species occurrence

The frequency of species occurrence between habitat states and the generalized linear model demonstrated that ecosystem engineering by beavers have a long-term effect on the frequency of mammalian species occurrence (Prediction 2, Fig. 4; see Supplementary Table S3). The three habitat states shared three species (moose, white-tailed deer, black bear), while ponds and wet meadows shared two more species (raccoon and coyote) (Fig. 4). Two species (muskrat and red fox) and one (bobcat) were only found at ponds and at wet meadows, respectively. Overall, more species were detected at ponds and wet meadows compared to dry meadows (Fig. 4). Coyotes were, on average, almost four times more frequent at wet meadows compared to ponds (0.023 ± 0.014, CI: 0.000–0.056 vs. 0.006 ± 0.004, CI: 0.000–0.015). However, the difference was not statistically significant according to confidence intervals overlap, while being all but completely absent from dry meadows. In contrast, white-tailed deer were almost four times more frequent at dry meadows compared to ponds (0.047 ± 0.012, CI: 0.024–0.070 vs. 0.012 ± 0.007, CI: 0.000–0.026), while black bears were, on average, only twice as frequent at dry vs. wet meadows (0.069 ± 0.018, CI: 0.033–0.105 vs. 0.028 ± 0.014, CI: 0.000–0.059). However, the difference was not statistically significant according to confidence intervals overlap.

**Figure 4:**
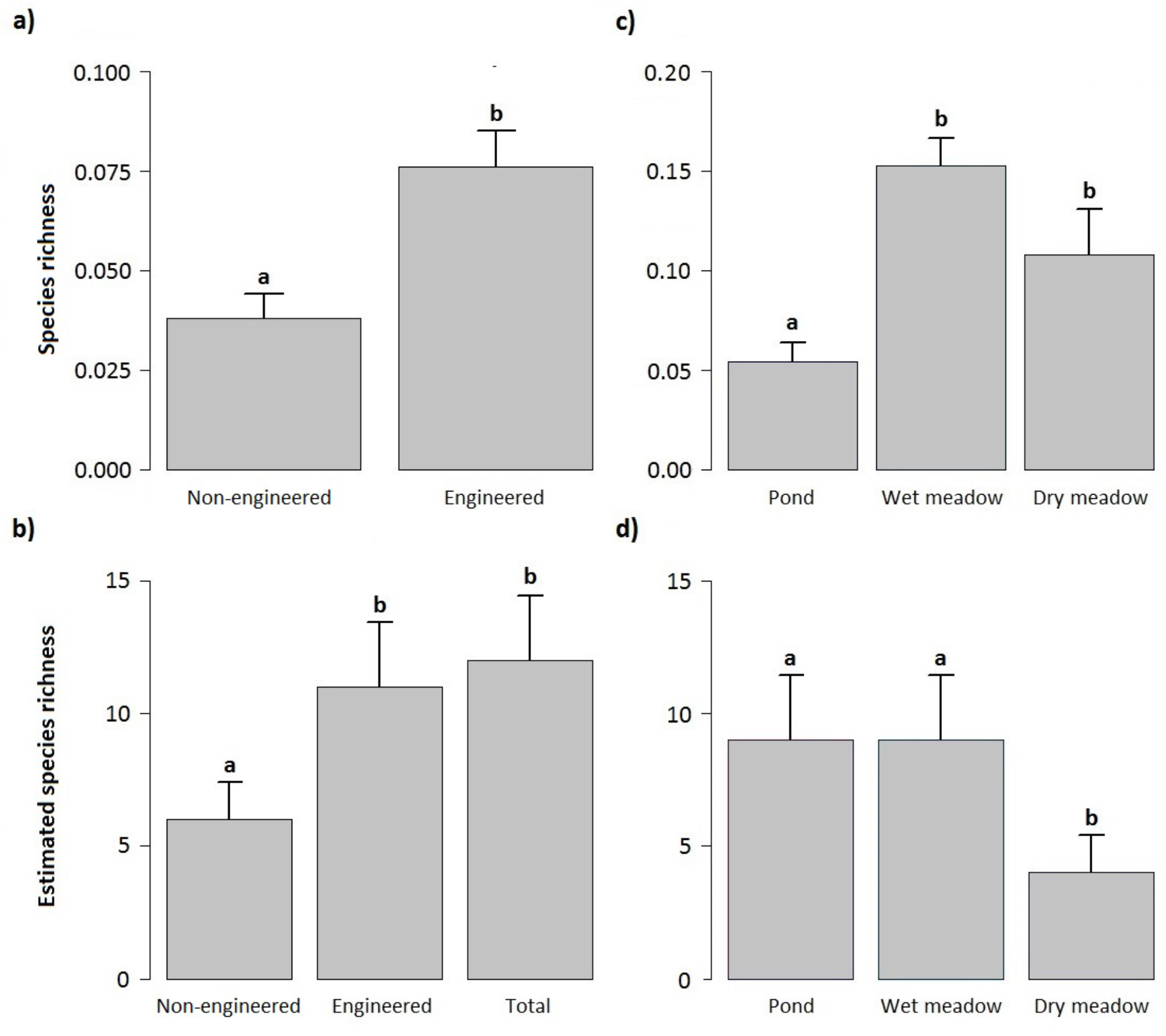
Frequency of species occurrence showing the effect of ecosystem engineering by beavers on the frequency of occurrence of individual species at engineered patches; **(a)** ponds (n= 9), **(b)** wet meadows (n= 2) and **(c)** dry meadows (n= 5), Kouchibouguac National Park of Canada, 2014–2015. The species frequencies were separated by year because two patches transitioned from one state to another from 2014 and 2015. We divided the total number of detections per day of each species by the number of camera trap days at each patch for each year and calculated the mean and the standard error for each patch types (n= 16 engineered patches per year). The detection period for each patch was over a 6-month period (i.e. 3 months per year). Error bars represent + 1 SE. Animal silhouettes were acquired from an open source (http://openclipart.org/).

### Habitat states and species composition

The mean Morisita-Horn similarity indices demonstrated that ecosystem engineering by beavers may also determine the species composition of mobile mammalian species in the long term (Prediction 2). Ponds and wet meadows were equally variable in species composition but more variable than dry meadows (0.39 ± 0.04, CI: 0.31–0.47 vs. 0.28 ± 0.10, CI: 0.08–0.47 vs. 0.66 ± 0.05, CI: 0.58–0.75; Fig. 2C). Ponds were, on average, no more similar to each other than they were to wet meadows (0.35 ± 0.04, CI: 0.27–0.43), but were less similar than they were to dry meadows (0.66 ± 0.05, CI: 0.57–0.75). Wet meadows were, on average, no more similar to each other than they were to dry meadows (0.39 ± 0.05, CI: 0.28–0.50). Comparisons of engineered patches from ponds, wet meadows, and dry meadows indicate an intermediate level of species overlap between the three habitat types. Accordingly, 62.5% of the 8 mammals’ species recorded at engineered patches in the survey were found in ponds and wet meadows, 42.9% in ponds and dry meadows, and 50% in wet meadows and dry meadows.

### Habitat states and species richness

Species richness of habitat states and the linear model demonstrated that ecosystem engineering by beavers has a long-term effect on the richness of mobile mammalian species at a local scale (Prediction 2). In meadows, species richness was at least twice higher compared to ponds (pond: 0.054 ± 0.009, CI: 0.037–0.071; wet meadows: 0.153 ± 0.014, CI: 0.126—0.180; dry meadows: 0.108 ± 0.022, CI: 0.065–0.150) (Fig. 3C, Table 3). The generalized linear model demonstrated that habitat states played a significant role in the observed species richness at ponds (p < 0. 001) and dry meadows (p = 0. 01), but not at wet meadows (p = 0. 139). Estimated species richness of habitat states and the linear model indicated that ecosystem engineering by beavers has a long-term effect on the overall richness of mobile mammalian species (Hyp. 2). Species richness estimates for all ponds and all wet meadows were identical (9 ± 2.45, CI: 7–17) but all higher than estimates for dry meadows (4 ± 1.41, CI: 3–9, Fig. 3D, Table 4). The linear model demonstrated that habitat states played a significant role in the observed estimated species richness at dry meadows (p = 0.028), but not at ponds (p = 0.127) and wet meadows (p = 0.362).

**Table 3:** Results of a generalized linear model showing the effect of habitat states (pond, wet meadow, and dry meadow) on species richness at 16 study sites, Kouchibouguac National Park of Canada, 2014–2015.

| Factors | Estimate | Standard error | *Z* | *P* |
| --- | --- | --- | --- | --- |
| Dry meadow | 0.108 | 0.015 | 7.054 | <0.0001 |
| Pond | -0.054 | 0.020 | -2.742 | 0.010 |
| Wet meadow | 0.045 | 0.030 | 1.521 | 0.139 |

Note: Pseudo $R^2 = 0.30$

**Table 4:**
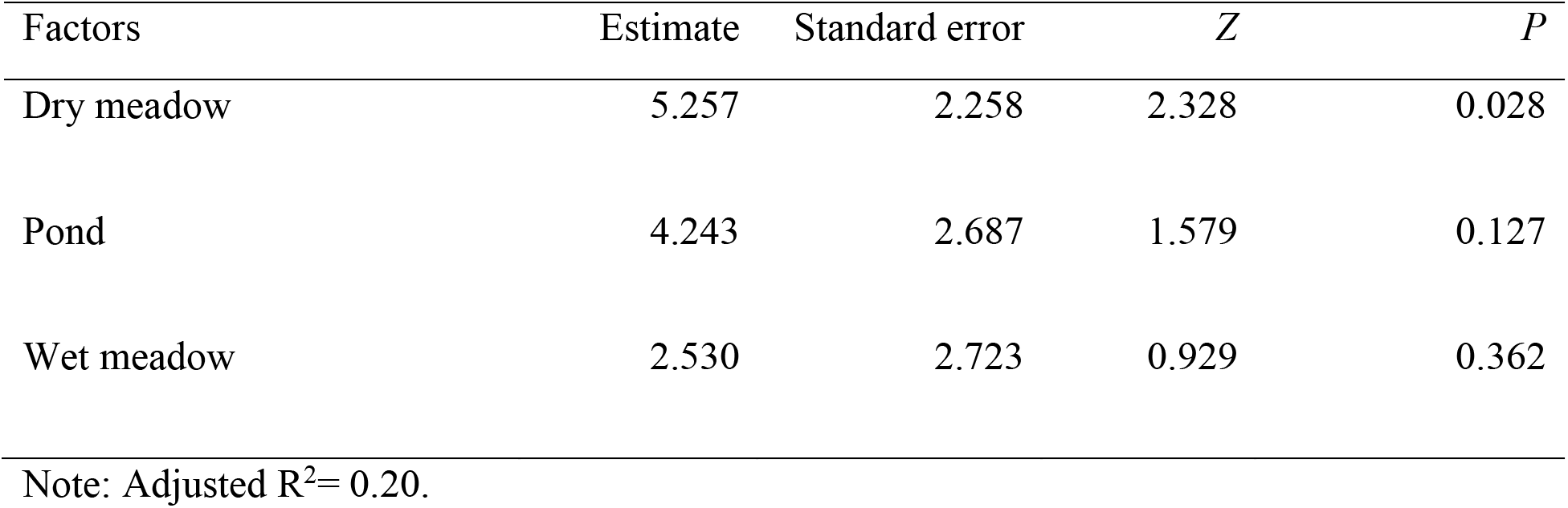
Results of a linear model showing the effect of habitat states (pond, wet meadow, and dry meadow) on species richness estimated from jackknife estimators at 16 study sites, Kouchibouguac National Park of Canada, 2014–2015.

| Factors | Estimate | Standard error | *Z* | *P* |
| --- | --- | --- | --- | --- |
| Dry meadow | 5.257 | 2.258 | 2.328 | 0.028 |
| Pond | 4.243 | 2.687 | 1.579 | 0.127 |
| Wet meadow | 2.530 | 2.723 | 0.929 | 0.362 |

Note: Adjusted $R^2 = 0.20$ .

## Discussion

The association between unique species and higher frequency of occurrence for most species at engineered patches support our hypothesis that ecosystem engineering by beavers increases species richness at local, but also at landscape scales for mobile mammalian species. The differences in frequency of species occurrence between the different types of engineered patches also support our second hypothesis that the state of habitats resulting from ecosystem engineering by beavers drives the diversity of mobile mammalian species. However, our data was limited with a few sites for certain habitat states, such as wet meadows, and therefore our findings will need to be confirmed by subsequent studies. Overall, such niche creation produced habitats heavily used across the landscape by those mobile mammalian species. Therefore, the impact of ecosystem engineering by beavers extends well beyond the habitats it creates, suggesting that beavers may play a considerable role in structuring terrestrial trophic interactions for species expressing a large range of dispersal and movement capabilities^13,22,28^.

Beaver-created habitats have been shown to benefit large mammals, especially species associated with wetlands such as muskrats^32,34^, otters^32,35,36,37^, and mink^32^. Such benefit could be exemplified for raccoons using beaver ponds in the summer months where aquatic vertebrates and invertebrates are abundant^32^. Beavers have also been shown to increase woody forage via their herbivory behavior, benefiting large herbivores, such as elk (*Cervus canadensis*)^38^, moose^39^ and white-tailed deer^32^, as well as smaller herbivores such as snowshoe hares^40^. Beaver meadows can also benefit large mammals, such as white-tailed deer and black bear^32^. Mesocarnivores have been documented using beaver lodges for shelter during winter (e.g. red fox)^41^ and as a breeding chamber during summer (e.g. Bobcat)^42^. Beavers are not a common prey for predators in our study area. For example, they only constitute a minor portion of the coyote’s diet in the park^43^ while black bears rarely feed on beavers^44^. Thus, the beaver population in our study area is not controlled by predators, but rather by the lack of their preferred food, trembling aspen^45^. This suggests that beavers themselves are not facilitating increased mesocarnivore density as prey. Therefore, the decrease of hare occurrences at engineered patches might be a cascading effect caused by an increase in mesocarnivores in those areas resulting from the ecosystem engineering aspects of beavers. A recent study, however, reported that beaver activity had no influence on the diversity and abundance of large mammals in the winter months^33^, suggesting that the impact of beaver-created habitats is not constant throughout the year.

We acknowledge that we did not detect all known large mammals’ species in our study area, with 50% of species detected (9 out of 18 mammalian species known to occur in our study area). However, the goal of our camera trap set-up was to detect a potential effect of ecosystem engineering by beavers on biodiversity and not explicitly to detect all known species of mammals in our study area. Accordingly, some species of mammals may not be detected even with a large survey effort of several thousands of camera trap days and our survey effort of 1,132 camera trap days is adequate to monitor mammalian species diversity^46^. Some of the species we missed were considered uncommon or linked to human infrastructure (e.g. American porcupine [*Erethizon dorsatum*] and stripped skunk [*Mephitis mephitis*])^47^. No camera-trapping system enables the detection of every mammalian species because terrestrial, arboreal and semi-aquatic species would each require a specific camera trap design^48^. For example, our camera trap design was aimed to detect terrestrial species while arboreal species, such as red squirrels (*Tamiasciurus hudsonicus*), would require a camera set up facing alongside of tree trunks. Aquatic species, such as river otters (e.g. *Lontra canadensis*) are rarely detected using camera traps based on passive infrared sensors such as those we used, because individuals recently out of the water have a low thermal signature and thus detection of such species require alternative trigger mechanisms (Lerone et al. 2015). Therefore, an important increase in our survey effort would not necessarily lead to the detection of such species, but could result in the detection of more medium-to large-sized terrestrial species. Detection rates were low for some species, namely predators (red fox, coyote and lynx) but their pattern of detection was robust, as they were all detected exclusively at beaver-engineered sites. In addition, to accurately estimate species occurrence, the distance between camera traps should be larger than the average home range of the species of interest to avoid spatial auto-correlation. Given our camera site density, we acknowledge that the occurrence of some species can be accurately estimated while others may not. Individuals of species with larger home ranges (e.g. moose, black bear, coyote) could be detected at more than one camera trap while individuals of species with smaller home ranges (e.g. raccoon, snowshoe hare) could be detected often at the same camera trap. Because our study design was not species-specific, we cannot accurately estimate the abundance of species. This implicates that there could be a high correlation between occurrence and abundance for species with small home ranges and little correlation for species with larger home ranges. We could not avoid this type of situation when the target measure is biodiversity, and the results of our analyses should be interpreted with such limitations in mind. Thus, for species with large home ranges, the response measure may reflect patterns of habitat selection whereas species with smaller home ranges, the response measure may better reflect local occupancy and abundance. To understand the effect of ecosystem engineering by beavers on the scales of movement across species, future studies could measure co-occurrence with the capture-mark-recapture method^70^.

The difference in species composition between patches in our study was mostly measured in terms of occurrence. The low species similarity among non-engineered patches was due to very few or no detection at most patches, while the intermediate similarity between engineered and non-engineered patches and the high similarity among engineered patches was due to the frequent detection of a few common mammalian species at most patches. Our results differ from those found for herbaceous plants^13^, which are adapted to particular light availability levels, soil moisture, and nutrient availability, for which changes in physical conditions following colonization by beavers^18^ triggered species turnovers. These differing results suggest that the effect of ecosystem engineering by beavers on species composition varies according to species life history traits. The differences that our study revealed in species composition within and between the different habitat states suggest that the effect of ecosystem engineering by beavers on mammalian species composition varies according to temporal changes in these engineered habitats. Ecosystem engineering by beavers and the habitat states of those engineered patches can then structure animal communities at local and landscape scales.

At the local scale, ecosystem engineering by beavers had a positive effect on species richness of mobile mammalian species. Our results differ from those found for stationary species, such as herbaceous plants^13^, and species which are relatively limited in their dispersal abilities, such as amphibians^22,24,25,26,27^. The latter demonstrated that ecosystem engineering by beavers had no predictable effect on species richness of those communities at the local scale. The local species richness of dispersal-limited species potentially depends on the number of species able to colonize the various engineered patches^50,51^. Therefore, it is difficult to predict higher or lower species richness at this scale for those communities. In contrast, the local species richness of mobile species seems to depend on the number of species present in the landscape (this study).

At the landscape scale, ecosystem engineering by beavers had a positive effect on species richness for mobile mammalian species. By attracting unique species, engineered patches increased the richness of mobile species at the landscape scale. Interestingly, similar results have been found for frogs, suggesting that such landscape scale effect could be true for a wider animal community^52,53^. This is similar to the results found for stationary species, but in a different manner. For instance, the difference in species composition of herbaceous plant species between engineered and non-engineered patches can increase the species richness at the landscape scale^13^. This occurred only when combining the species found at both types of patches. Furthermore, the association of most mammalian species to ponds and wet meadows suggests that the presence of wetlands and not just open habitats and rivers might be important to attract such diversity of mobile species across the landscape. Nonetheless, the association of some mammalian species to different states of habitats created by beavers suggests that the impact of such ecological engineers extends well beyond the initial habitat flooding and creates long-term dynamics of habitat use. This is not surprising considering that parts of tributaries with a history of beaver occurrence are more geomorphologically complex, biologically diverse, and productive than the surrounding landscape^54^. We know that some ponds or beaver meadows can be ≥40 years old^19,20,45^. While the introduction of beavers to new habitats (e.g. in Argentina)^55^ can start a long ecological succession, the species’ imprint may last despite population declines^45^. In a management context, this underlines the potential repercussions of niche creation at a large spatial scale and over a long ecological period.

This study is a step towards understanding the larger influence of ecosystem engineering on important components of ecosystems. It illustrated such a spillover effect by focusing on key species of the boreal biome, possibly modifying biodiversity at multiple scales well beyond the engineered structures. Species life history traits and study scale are thus both predictors of the impact of ecosystem engineers on communities. The imprint of ecological engineering and niche creation on the functioning of many ecosystems may then be examined through the lenses of ecological succession and dispersal capabilities.

## Methods

### Study area

The study was conducted at Kouchibouguac National Park of Canada, an area of 238.8 km^2^ on the east coast of New Brunswick (46 °42’ to 46 °57’ N, 64 °47’ to 65 °02’ W)^56^. Founded in 1969, the park is located within the Maritime Lowlands Ecoregion, hosting forests of red spruce (*Picea rubens*), black spruce (*Picea mariana*), balsam fir (*Abies balsamea*), red maple (*Acer rubrum*), eastern hemlock (*Tsuga canadensis*), as well as white pine (*Pinus strobus*) and jack pine (*Pinus banksiana*)^57^. This area has a humid continental climate with important maritime influences^58^. The flat topography and the low permeability of soils support numerous peatlands and swamps^57,58^. The park is dominated by eight major watersheds and two large peatlands^56^. Forests cover more than half of the park (54.7%)^56^ with numerous mixed and coniferous stands, and a few deciduous stands^58,59^.

The park’s beaver population is protected from fur trapping since the establishment of the park in 1969, although some trapping may have occurred in the few years following park’s creation. This enabled a population dynamic without harvest, leading to a spatial mosaic of heterogeneous habitats in different states across the landscape. Typically, sites occupied by beavers are ponds that from as a result of flooding caused by the construction and maintenance of a dam made of tree branches and mud. Once abandoned by beavers, they transition into a meadow dominated by non-woody vegetation or bushes. Such sites may become dry habitat with the original stream running through it (dry meadow) or may have a residual pond and marshy area (wet meadow). In 2014, we did an on-foot census of first to third order streams and found 271 sites with a history of beaver occurrence^45^. The census revealed that approximately 28% of the entire riparian zone had a history of beaver occurrence. Looking back to historical records, the natives of Acadia began to trade fur, particularly beaver^60^, with the French in the early 1600s^61,62,63^ and kept trading well after the final conquest of New France in 1760, and until the late 1890s^64,65^. The earliest documented instances of trapping in this region occurred nearly a century after records stating that beavers damming watersheds constituted major barriers for early explorers using canoes^66^. Therefore, the beaver-mediated mosaic of habitats that currently exists in the park is likely representative of conditions found before the advent of considerable anthropogenic pressures such as forest exploitation, which became rampant in this region during the 19^th^ century^67,68^.

### Field methods

To evaluate the spatial effect of ecosystem engineering by beavers on the diversity of mobile mammalian species, we surveyed 16 sites encompassing multiple watersheds and patches with and without a history of beaver occurrence (Hyp. 1). We performed our study during the summers of 2014 and 2015 using camera-trapping techniques (Fig. 5). Therefore, we collected data inside and outside the beaver’s foraging extent on land (i.e. 100 m from the edge of the pond or meadow)^59,69^ at each study site. One camera trap was set up at the beaver infrastructure (i.e. engineered patches), and the other at least 400–500 m away (i.e. non-engineered patches). At each study site, camera traps at both engineered and non-engineered patches were installed near tributaries, at similar distances from human perturbations (e.g. roads) and main watersheds, and at places with similar plant community composition. Because the study area was saturated by patches with a history of beaver occurrence, we could not select non-engineered sites further upstream or downstream in the same tributaries as the engineered patches. Non-engineered sites were therefore located in the vicinity of tributaries that had no history of beaver occurrence^45^ (Fig. 5). To quantify the long-term impact of ecosystem engineering by beavers on the diversity of mobile mammalian species (Hyp. 2), the 16-engineered patches selected for our study included three habitat states present in our study area, i.e. ponds (n= 9), wet meadows (n= 2), and dry meadows (n= 5). Wet meadows were not as common as the other habitat states in our study area, however preliminary analysis showed interesting biodiversity responses to this habitat state and were therefore kept in our analysis despite their low number of samples. Overall, our camera density is ca 0.016 per km^2^ (ca 1 camera for each 15km^2^ of the 238.8 km^2^ park).

**Figure 5:**
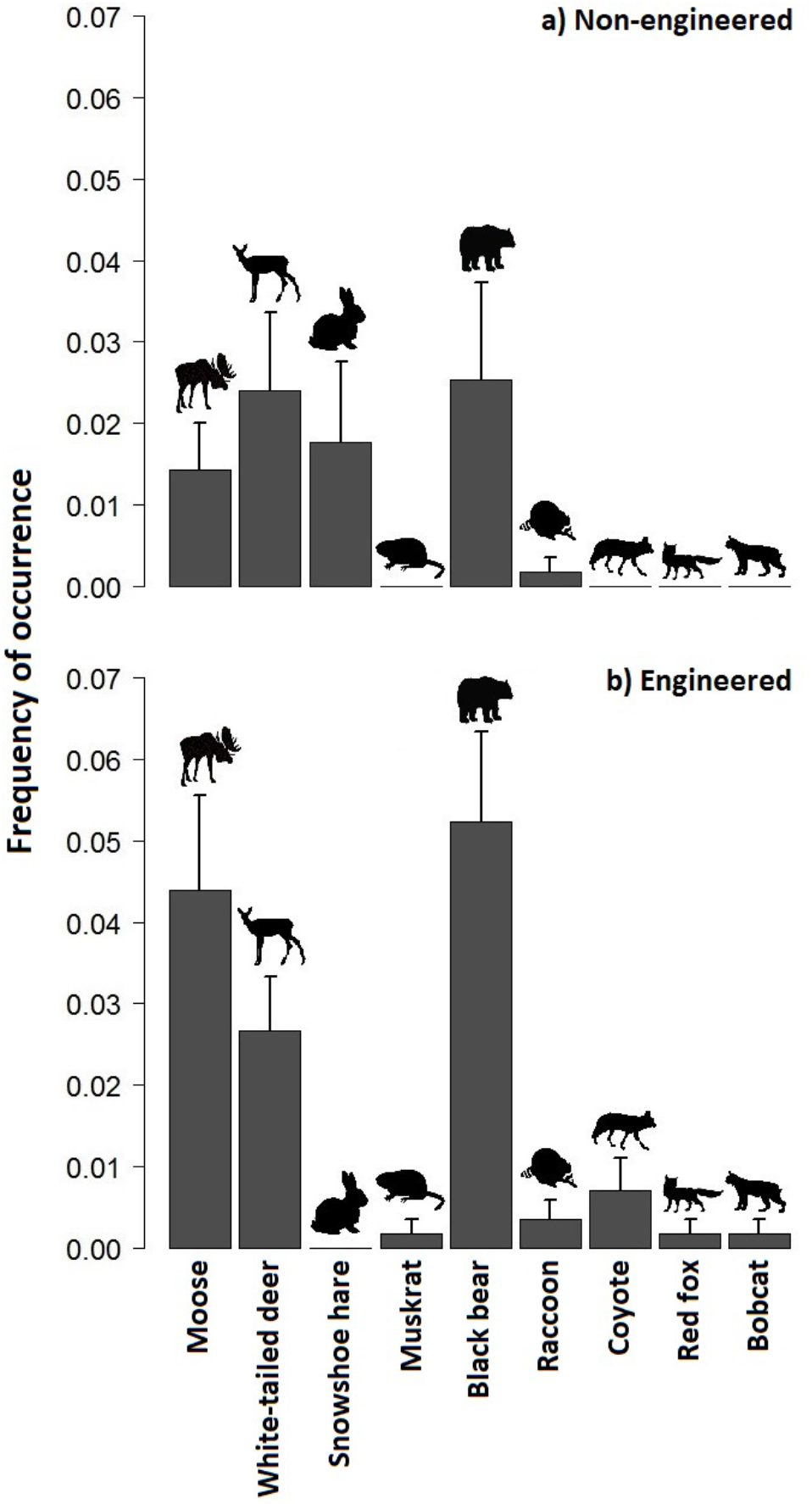
Study area at Kouchibouguac National Park of Canada, New Brunswick, Canada, showing our 16 study sites sampled in 2014–2015. Each study site encompasses an engineered and a non-engineered patch. The engineered patches were within the beaver’s home range (at the pond’s edge) and the non-engineered patches were 400–500 meters away from engineered patches. The figure was produced using a modified version of a New Brunswick province map from the Open Government License — Canada (Copyright: Her Majesty the Queen in Right of Canada, Natural Resources Canada, 2002) and using QGIS software 2.4.0^84^.

We used a passive method of detection (i.e. no lures) to detect mammals with our camera traps in a similar manner across our study area and camera traps were set up to maximize detection of a wide variety of mammal species and were not species specific. Cameras were in operation 24 hours daily to maximize chances of detection. We rotated 8 devices among study sites on a weekly schedule over a period of 12 weeks; from June 4^th^ in 2014 and from May 26^th^ in 2015. This camera rotation regime enabled us to increase the number of study sites despite a limited number of cameras. Cameras are a common limiting factor in camera trap studies because of their cost and often require a rotation regime to reach a higher number of sampling sites^70^. Each week, we installed a device at the engineered and the non-engineered patch for four study sites. Camera traps were set up in a manner where the visibility was similar between each patch. They were facing alongside the edge of engineered or non-engineered patches without water body in the camera frame to detect mammals coming to these areas. Cameras were set at natural openings and non-woody vegetation was cleared in a standardized manner at areas without openings. At each patch, camera traps were always set up at the same location, facing the same direction, and at the same height and angle. Cameras should be set up at the same height as the core mass of the target species^71^, yet our camera traps were not species specific. Therefore, cameras were set at chest height to detect larger species with a facing downward angle, which is preferred for the detectability of smaller species.

Each series of pictures featuring an animal (from its entry until its departure from the camera frame) was counted as a single detection within a 24-hour period. The camera models were HC600 Hyperfire H. O. Covert IR (Reconyx Inc.) and Spypoints (IR-5, IR-6, BF-7, and Tiny-W). The Reconyx Inc. cameras were set to take three pictures per trigger with one-second delay between pictures and no delay between triggers; the rest of the settings were by default (e.g. high-resolution pictures). The same settings applied for the Spypoint cameras, except that four pictures were taken per trigger, with a one-minute delay between triggers. This setting was the closest possible match with the Reconyx cameras’ settings. The simultaneous monitoring of both the engineered and the non-engineered patch of a given site was always done with the same camera type. In 2015, we used Reconyx Inc. cameras to collect data exclusively, and used the other camera types in a comparative experiment to determine differences in performance to help interpret the 2014 dataset. We installed Spypoint cameras in tandem to Reconyx Inc. cameras to compare the number of animal detections between them. We did not find any difference between camera brands (χ ^2^ = 2.24, *d*.*l*. = 1, *n* = 28, *p* = 0.13).

### Data analyses

To evaluate the potential effect of ecosystem engineering by beavers on the frequency of species occurrence (Prediction 1.1), we calculated the frequency of occurrence of each mammalian species for engineered and non-engineered patches. This was obtained by dividing the total number of animal detections of each species by the number of camera trap days at each patch^72^ to account for the differences in the number of camera trap days at each patch, and calculated the mean, the standard error, and the confidence intervals for both patch types. We then performed a generalized linear model (Poisson distribution) to determine whether the patch type (engineered vs. non-engineered) played a significant role in the observed frequency of occurrence of each species.

To evaluate the potential effect of ecosystem engineering by beavers on mammalian species diversity, we started by comparing the species composition between engineered and non-engineered patches using two techniques (Prediction 1.2). First, we performed a non-metric multidimensional scaling ordination based on species presence-absence data for each patch to portray the degree of species overlap between engineered and non-engineered patches. Second, we calculated Morisita-Horn similarity indices for pairwise comparisons of patches based on the frequency of occurrence for species found within each patch. This index measures the similarity between two communities by comparing the species composition within and between patches type (engineered and non-engineered) and can be interpreted as a probability^73^.

We compared the mammalian species richness between engineered and non-engineered patches at local and landscape scales using four techniques (Prediction 1.2). First, we calculated the species richness at individual engineered and non-engineered patches to evaluate the potential effect of ecosystem engineering by beavers on species richness at the local scale. The species richness was obtained by dividing the number of species by the number of camera trap days at each patch to account for differences in the number of camera trap days at each patch, and then by calculating the mean, the standard error, and the confidence intervals for both patch types. Second, to quantify the potential effect of engineered and non-engineered patches on species richness, we performed a generalized linear mixed model with a Poisson distribution, using study sites as random factor. Third, we used estimates of species richness to calculate the potential effect of ecosystem engineering by beavers on species richness at the landscape scale by estimating species richness of a landscape without beaver-modified habitats (i.e. all non-engineered patches), and then of an equally sized area with beaver-modified habitats (i.e. all engineered patches). This provided an estimate of the contribution of engineered and non-engineered areas to the total species richness of the riparian zone (i.e. all engineered and non-engineered patches combined). Estimates were calculated based on the frequency of occurrence of each species for all engineered patches, all non-engineered patches and all study sites (engineered and non-engineered patches). We used a first order Jackknife estimator to do so, taking advantage of this resampling method to minimize bias in our estimates. Fourth, we applied a linear model to determine the potential effect of all engineered patches, all non-engineered patches, and all study sites on the overall species richness because jackknife estimators were drawn from a normal distribution.

Similar analyses were used to evaluate the potential effect of the habitat states (ponds, wet meadows and dry meadows) of engineered patches on the frequency of occurrence of individual mammalian species, species composition and richness (Prediction 2). To determine whether habitat states played a significant role in the frequency of species occurrence, we calculated the frequency of occurrence of each species and performed a generalized linear model (Poisson distribution). We then calculated Morisita-Horn similarity indices to evaluate the effect of species composition. We calculated the species richness and performed a generalized linear model (normal distribution) to determine whether the habitat states played a significant role in the species richness at the local scale. Finally, we calculated jackknife estimators and performed a linear model to determine whether the habitat states played a significant role in the overall species richness.

Rotation days of camera traps were excluded from analyses because our presence may have influenced the behavior of some animals. Central tendencies are expressed as mean ± SE or with 95% confidence intervals. When performing linear models, significance tests and their associate p-values were used to determine statistical significance. When comparing central tendencies, estimates with confidence intervals were used to determine statistical significance because they are equivalent to a significance test but convey more information^74^.

We performed all statistical analyses using R version 3.2.2^75^. We used the packages MASS^76^, vegan^77^ and BiodiversityR^78^ to perform non-metric multidimensional scaling, vegetarian^79^ to perform Morisita-Horn similarity indices, R2admb^80^ and glmmADMB^81^ to perform generalized linear mixed models, bbmle^82^ for model comparisons and SPECIES^83^ to perform jackknife estimations. We verified all assumptions linked to the use of our statistical models (e.g. heteroscedasticity, normality, Poisson distributions).

## Acknowledgements

We thank N. Tran and J. Dubé for their help with fieldwork. This work is part of an MSc project in the Canada Research Chair in Polar and Boreal Ecology at Université de Moncton, which benefited from early comments by G. Moreau and M.-A. Villard. We thank S. Craik for his input during the revision of this work. We are grateful for the logistical support of Kouchibouguac National Park of Canada and the Université de Moncton. Funding was provided by: the Canada Research Chair Program, the Natural Sciences and Engineering Research Council of Canada, the Canadian Foundation for Innovation, the New Brunswick Foundation for Innovation, the New Brunswick Wildlife Trust Fund, the Canadian Heritage, and the Université de Moncton.

## Author contributions

NL, DG, and LG conceived and designed the study. DG, LG, and NL ran the fieldwork. DG, LG, and NL performed the analyses. All authors participated in the writing.

## Additional Information

We do not have any competing financial and non-financial interests.

