## Supplementary Material for "Spatiotemporal changes in biodiversity by ecosystem engineers: how beavers structure the richness of large mammals"

[1] Canada Research Chair in Polar and Boreal Ecology and Centre for Northern Studies, Université de Moncton, 18 Antonine-Maillet, Moncton, New Brunswick, E1A 3E9, Canada.

+ Senior author

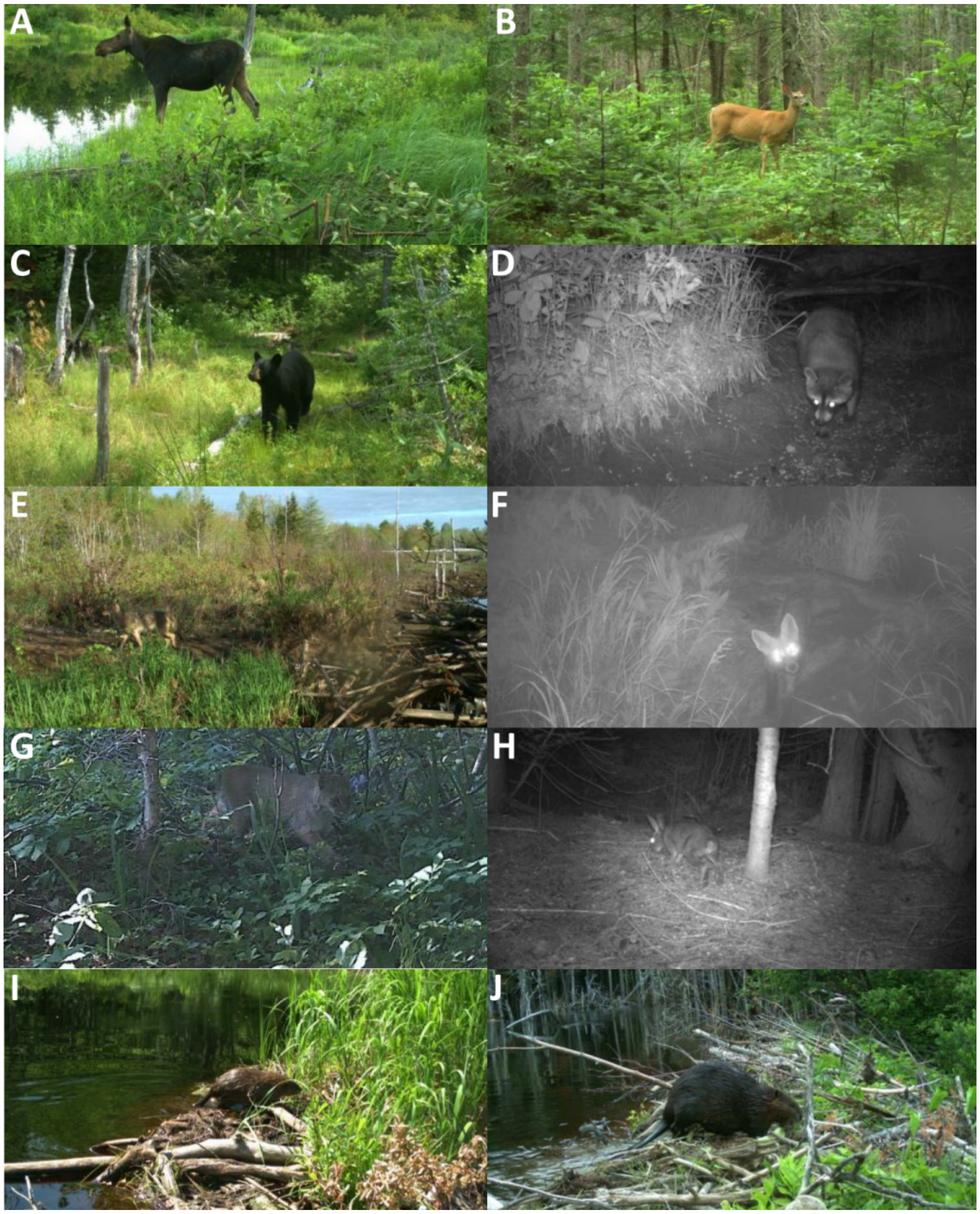


Supplementary Figure S1: The beaver and the 9 species of mammals captured by camera traps during the summers of 2014 and 2015 in 16 study sites, Kouchibouguac National Park of Canada. (**A**) Moose, (**B**) white-tailed deer, (**C**) black bear, (**D**) raccoon, (**E**) coyote, (**F**) red fox (**G**) bobcat, (**H**) snowshoe hare, (**I**) muskrat, and (**J**) beaver. Some animal pictures were cropped and/or taken from a collection of photographs from another research project using camera traps in our study area to better illustrate our focal species.

Supplementary Tables

Supplementary Table S1: Regression coefficients of a Generalized Linear Model showing the effect of patch type (engineered vs. non-engineered) on the frequency of occurrence of individual species (species identity as factor and frequency of occurrence as response variable) at 16 sites, Kouchibouguac National Park of Canada, 2014–2015.

| Factors | Estimate | Standard error | *Z* | *P* |
|---|---|---|---|---|
| Intercept: Moose*Engineered patch | 0.044 | 0.006 | 7.055 | <0.0001 |
| White-tailed deer | -0.017 | 0.009 | -1.964 | 0.051 |
| Black bear | 0.008 | 0.009 | 0.939 | 0.348 |
| Raccoon | -0.041 | 0.009 | -4.595 | <0.0001 |
| Coyote | -0.037 | 0.009 | -4.200 | <0.0001 |
| Snowshoe hare | -0.042 | 0.009 | -4.792 | <0.0001 |
| Bobcat | -0.044 | 0.009 | -4.989 | <0.0001 |
| Red fox | -0.042 | 0.009 | -4.792 | <0.0001 |
| Muskrat | -0.042 | 0.009 | -4.792 | <0.0001 |
| Non-engineered patch | -0.030 | 0.009 | -3.368 | 0.001 |
| White-tailed deer x non-engineered patch | 0.027 | 0.012 | 2.170 | 0.031 |
| Black bear x non-engineered patch | 0.003 | 0.012 | 0.224 | 0.823 |

| | | | | |
|---|---|---|---|---|
| Raccoon x non-engineered patch | 0.028 | 0.012 | 2.242 | 0.026 |
| Coyote x non-engineered patch | 0.023 | 0.012 | 1.824 | 0.069 |
| Snowshoe hare x non-engineered patch | 0.028 | 0.012 | 2.242 | 0.026 |
| Bobcat x non-engineered patch | 0.047 | 0.012 | 3.802 | <0.0001 |
| Red fox x non-engineered patch | 0.028 | 0.012 | 2.242 | 0.026 |
| Muskrat x non-engineered patch | 0.028 | 0.012 | 2.242 | 0.026 |

Note: Pseudo $R^2$= 0.30.

Supplementary Table S2: Regression results of a Generalized Linear Model showing the effect of habitat states (pond, wet meadow, and dry meadow) on the frequency of occurrence of individual species (species identity as factor and frequency of occurrence as response variable) at 16 engineered patches, Kouchibouguac National Park of Canada, 2014–2015.

| Factors | Estimate | Standard error | *Z* | *P* |
|---|---:|---:|---:|---:|
| Moose x Pond | 0.028 | 0.008 | 3.472 | 0.001 |
| White-tailed deer | -0.016 | 0.012 | -1.385 | 0.167 |
| Black bear | 0.022 | 0.012 | 1.885 | 0.061 |
| Raccoon | -0.025 | 0.012 | -2.188 | 0.030 |
| Coyote | -0.022 | 0.012 | -1.920 | 0.056 |
| Snowshoe hare | -0.028 | 0.012 | -2.455 | 0.015 |
| Bobcat | -0.028 | 0.012 | -2.455 | 0.015 |
| Red fox | -0.025 | 0.012 | -2.188 | 0.030 |
| Muskrat | -0.028 | 0.012 | -2.455 | 0.015 |
| Wet meadow | 0.111 | 0.019 | 5.775 | <0.0001 |
| Dry meadow | 0.007 | 0.014 | 0.526 | 0.600 |
| White-tailed deer x Wet meadow | -0.081 | 0.027 | -3.000 | 0.003 |
| Black bear x Wet meadow | -0.147 | 0.027 | -5. 421 | <0.0001 |

| | | | | |
|---|---|---|---|---|
| Raccoon x Wet meadow | -0.100 | 0.027 | -3.684 | 0.003 |
| Coyote x Wet meadow | -0.089 | 0.027 | -3.285 | 0.001 |
| Snowshoe hare x Wet meadow | -0.097 | 0.027 | -3.570 | <0.0001 |
| Bobcat x Wet meadow | -0.111 | 0.027 | -4.083 | <0.0001 |
| Red fox x Wet meadow | -0.114 | 0.027 | -4.198 | <0.0001 |
| Muskrat x Wet meadow | -0.097 | 0.027 | -3.570 | <0.0001 |
| White-tailed deer x Dry meadow | 0.027 | 0.019 | 1.419 | 0.157 |
| Black bear x Dry meadow | 0.012 | 0.019 | 0.614 | 0.540 |
| Raccoon x Dry meadow | -0.010 | 0.019 | -0.532 | 0.595 |
| Coyote x Dry meadow | -0.013 | 0.019 | -0.692 | 0.490 |
| Snowshoe hare x Dry meadow | -0.007 | 0.019 | -0.372 | 0.710 |
| Bobcat x Dry meadow | -0.007 | 0.019 | -0.372 | 0.710 |
| Red fox x Dry meadow | -0.010 | 0.019 | -0.532 | 0.595 |
| Muskrat x Dry meadow | -0.007 | 0.019 | -0.372 | 0.710 |

Note: Pseudo $R^2$= 0.35.
